# Sequence analysis of Indian SARS-CoV-2 isolates shows a stronger interaction of mutated receptor binding domain with ACE2 receptor

**DOI:** 10.1101/2020.08.28.271601

**Authors:** Pujarini Dash, Jyotirmayee Turuk, Santosh Ku. Behera, Subrata Ku. Palo, Sunil K. Raghav, Arup Ghosh, Jyotsnamayee Sabat, Sonalika Rath, Subhra Subhadra, Debadutta Bhattacharya, Srikant Kanungo, Jayasingh Kshatri, Bijaya kumar Mishra, Saroj Dash, Namita Mahapatra, Ajay Parida, Sanghamitra Pati

**Affiliations:** Regional Medical Research Centre, Indian Council of Medical Research, Bhubaneswar, Odisha, India; Institute of Life Sciences, Nalco square, Chandrasekharpur, Bhubaneswar, and Odisha, India

**Author notes:** ***Author for correspondence***: Dr. Jyotirmayee Turuk, Dr. Sunil K. Raghav, ICMR–Regional Medical Research Centre, Chandrasekharpur, Bhubaneswar – 751023, Odisha, India.

**Keywords:** SARS-COV-2, Indian isolates, Spike, RBD, ACE2

## Abstract

SARS-CoV-2 is a RNA Coronavirus responsible for the pandemic of the Severe Acute Respiratory Syndrome (COVID-19). It has affected the whole world including Odisha, a state in eastern India. Many people migrated in the state from different countries as well as states during this SARS-CoV-2 pandemic. As per the protocol laid by ICMR and Health & Family welfare of India, all the suspected cases were tested for SARS-CoV-2 infection. The aim of this study was to analyze the RNA binding domain (RBD) sequence of spike protein from the isolates collected from the throat swab samples of COVID-19 positive cases and further to assess the RBD affinity with ACE2 of different species including human.

Whole genome sequencing for 35 clinical SARS-CoV-2 isolates from COVID-19 positive patients was performed using ARTIC amplicon based sequencing. Sequence analysis and phylogenetic analysis was carried out for the Spike and RBD region of all isolates. The interaction between the RBD and ACE2 receptor of five different species was also analysed.

Except three isolates, spike region of 32 isolates showed one/multiple alterations in nucleotide bases in comparison to the Wuhan reference strain. One of the identified mutation at 1204 (Ref A, RMRC 22 C) in the RBD of spike protein was identified which depicted a stronger binding affinity with human ACE2 receptor compared to the wild type RBD. Furthermore, RBDs of all the Indian isolates are capable of binding to ACE2 of human, bat, hamster and pangolin.

As mutated RBD showed stronger interaction with human ACE2, it could potentially result in higher infectivity. The study shows that RBDs of all the studied isolates have binding affinity for all the five species, which suggests that the virus can infect a wide variety of animals which could also act as natural reservoir for SARS-CoV-2.

## 1. Introduction

Corona viruses have been studied for more than fifty years and are known to infect multiple animal species including human. Though their pathogenesis and mechanism of replication has already been well described due to previous outbreaks (SARS-CoV in China, 2003 and MERS-CoV in Saudi Arabia, 2012), the current pandemic has propelled the whole world to research and investigate deeper into the pathogenesis of this virus. Coronaviruses belong to a diverse virus family which consists of four genera: Alpha, Beta, Gamma and Delta corona viruses^1^. According to the nucleic acid sequence similarity, SARS-CoV-2 or COVID-19 is a beta corona virus. The spike glycoprotein of corona viruses facilitates its entry into their host cell and also gives the virus a crown like structure on its surface. Binding of pathogenic particles of COVID-19 with host cell receptors remains the crucial step for initiation of infection. The key factor lies in receptor recognition and establishment of infection in cells or tissues. Besides, the ability of the virus to bind to the specific receptor of other host species is also an essential requirement for transmission across different species^2^. It is known that SARS-CoV interacts with human angiotensin converting enzyme 2 (ACE2) for their entry and after COVID-19 outbreak, researchers found that SARS-CoV-2 also interacts with ACE2 for the viral entry into host cells^3,4^. SARS-CoV-2 spike (S) protein gets cleaved by host proteases into S1 and S2 domain which mediates receptor recognition and membrane fusion respectively^5^. Wang et al. (2020) reported that, the S1 domain of S protein of SARS-CoV-2 contains a region called receptor binding domain (RBD) which makes a complex with human ACE2 and facilitates viral entry^6^. However, emerging mutations in SARS-CoV-2 genome might alter the process of infection transmission, replication and potential of viral attachment with ACE2. According to epidemiological data, COVID-19 has an origin from bats in Wuhan, China and then spread to other parts after its zoonotic transmission via the Malayan Pangolins^7,8^. In a country like India having a diversified geographical distribution, it is important to understand the pathogenesis of different strains of SARS-CoV-2 isolated from different parts of the country. As interaction with ACE2 is the main pathway of entry of this virus into its host, knowledge on the RBD binding affinity of different Indian SARS-CoV-2 isolates with the ACE2 of their natural reservoirs including human is highly essential. However, this has remained a grey area with a little information available. In the current study, we have analysed the RBD sequences of spike protein from different isolates of SARS-CoV-2 from COVID-19 patients of Odisha. Further we analysed the binding affinity of RBDs with the ACE2 of different probable natural hosts of corona virus: bat, pangolin and hamster including human. The detected mutation in the RBD region of one isolate shows stronger binding affinity with human ACE2 than the wild type RBD, providing important information regarding its virulence as well as drug targeting.

## 2. Material & Methods

### Sequencing of different Indian isolates of SARS-CoV-2 from throat swab samples

The current study was a part of whole genome sequencing study carried under the Odisha Study group constituting different Government organisations jointly by Regional Medical Research Centre (RMRC), Bhubaneswar and Institute of Life Sciences, Bhubaneswar. As ICMR-RMRC, Bhubaneswar is a govt. authorised testing laboratory for COVID-19 testing, we received throat swab samples of suspected cases from different hospitals of Odisha. For COVID-19 diagnosis, viral RNA isolation was carried out from all samples using QIAmp Viral RNA Mini Kit followed by qPCR (Taqpath™ 1-step Master Mix, ThermoFisher Scientific). The libraries were prepared for WGS using ACTIC amplicon based sequencing kits from Qiagen as per manufacturer recommended protocol. Out of all positive samples, whole genome sequencing of 35 isolates from different COVID-19 patients with different travel history, was carried out using Illumina platform. The detailed information and method have been described by Raghav et al., 2020^9^

### Sequence alignment of RMRC spike genes with reference sequence

For sequence alignment, full length genome sequence of SARS-CoV-2 isolate of Wuhan-Hu-1 (Accession no.NC_045512) was downloaded from NCBI database and used as reference sequence for all further analysis. Alignment of all 35 RMRC spike nucleotide sequences with the reference genome was carried out using BLASTN (align two/more sequences).

### Identification and phylogenetic analysis of RBD of RMRC SARS-CoV-2 isolates

From the data available for reference strain, we retrieved the information of coding region of spike RBD domain sequence and all RMRC spike sequences were aligned with the reference sequence to identify the respective RBD mutations using BLASTN. Multiple sequence alignment of all 35 RBDs (amino acid sequences) was carried out using Clustal X tool. Mutations specific to RMRC isolates were identified by comparing the RBD coding regions with the reference strain. A phylogenetic tree was generated using the MEGA software version 6 with 1000 bootstrap replications as instructed in MEGA software.

### Sequence and 3D structure analysis of ACE2 receptor

From the phylogenetic tree analysis, four RMRC RBD sequences were selected from four random clusters for further investigation of their interactions with ACE2 receptor of probable natural hosts of SARS-CoV-2. Hamsters were reported to be the best suitable animal model to carry out SARS-CoV-2 related experiments; therefore it was essential to understand the interaction of its ACE2 receptor with isolated Indian SARS-CoV-2 RBDs^10^. For the interaction study, hamster (*Mesocricetus auratus*) (UniprotKB-C7ECV1, 785 amino acids), pangolin (*Manis javanica)* (NCBIXP_017505752, 805 amino acids), Chinese bat (*Rhinolophus sinicus*) (Uniprot E2DHI7, 805 amino acids), Indian bat (*Cynopterus sphinx)* (Uniprot QZF77831, 807 amino acids) (probable natural hosts for SARS-CoV-2) and human (*Homo Sapiens*) ACE2 (UniprotKB Q9BYF1, 805 amino acids) sequences were included. Before inception of structure prediction, the ACE2 sequences from pangolin, hamster, Chinese and Indian bat were aligned with the human counterpart to identify the percentage of similarity and dissimilarity between these sequences.

The experimental 3D structure of human ACE2 was retrieved from RCSB PDB (PDB ID-6M0J) with a resolution of 2.45 Å positioning from 19-615 amino acids. Protein Data Bank (PDB) did not provide any experimental structure of ACE2 receptors of pangolin, hamster, Chinese and Indian bat, which has prompted us to predict their three dimensional (3D) structure through homology modelling using Modeller v 9.19 tool followed by structure validation. Suitable templates were identified for 3D model building of ACE2 using BLASTp^11^ search against PDB (Protein Data Bank) database. The templates with PDB ID: 1R42, 6CS2, 6LZG, 3SCI, and 2AJF, were found to be the better homologs for target-template alignment and modelled 3D structure prediction. Based on the optimized target-template alignment, Modeller v9.19^12^ facilitated in model developments, the models with lowest Discrete Optimized Protein Energy (DOPE) score were retained for further structural refinement. Side chain optimization was performed using WHATIF^13^ and GalaxyRefine^14^ tool. The optimized models of ACE2 were finalized based on its overall quality and stereo-chemical geometry and energy. The geometry of the predicted model was evaluated using PROCHECK^15^ and Ramachandran analysis^16^. ERRAT^17^ programme was used to calculate the accuracy of the non-bonded atoms for the predicted model. Verify 3D^18^ tool was used to evaluate the compatibility of the 3D model with its own amino acid sequence by assigning a structural class based on its location, environment and comparing the results to good quality structures. The structure was uploaded in Qualitative Model Energy Analysis (QMEAN) server (bencrept), for resolving the model quality. The energy potential of the predicted model was calculated using ProSA-web server^19^.

### *In silico* translation of RBD sequences and their interaction with ACE2 receptor of different natural reservoirs

Four nucleotide sequences of RMRC RBDs were selected and translated to protein sequence using EMBOSS Transeq tool of European Bioinformatics Institute (EBI). The 3-D structure of reference human RBD (Uniprot ID: PODTC2, 333-526 amino acids of the spike protein, PDB ID-6M0J) was considered as wild type. The experimental structure of the wild RBD was mutated at the position I402L using Discovery Studio visualizer (4.1) for attaining a mutant RMRC22 RBD as required for further computational analysis. Finally, study of protein-protein interaction (PPI) between wild type/mutant RBDs and ACE2 receptors form different organisms was carried out using online HawkDock server which is a powerful tool to predict the binding structures and identify the key residues of PPIs^20^.

## 3. Results

### Sequence information and analysis of spike gene of Indian SARS-CoV-2 isolates

All the SARS Co-V-2 isolates included in the study had a travel history from either outside the country or from other states of India. According to the travel history, out of 35 isolates, one had a foreign travel history, 11 were Nizamuddin (cluster detected from New Delhi, India during April, 2020) returned and rest 23 migrated from Surat, Gujarat, India. The detail of the demographic and clinical status of the patients included in the study has been described in one of our unpublished reports (Turuk et al). We did not record any death case among the patients whose samples were included in the current study.

. In the current study, the spike region was identified at the position from 21563-25384 of the whole genome and consists of 3822 nucleotides. The BLAST alignment analysis of RMRC spike nucleotide sequences alignment analysis showed that, three spike sequences (RMRC 104, 157, 158) were 100 % identical to the Wuhan reference spike whereas all other RMRC spikes shared 99 % identity with one or multiple altered bases at different positions (Table 1).

**Table 1:** Patient information and sequence identity analysis of all the RMRC SARS-CoV-2 isolates.

| Sl. No. | Sample ID | RMRC Spike nucleotide seq. identity with reference (%) | RMRC RBD nucleotide seq. identity with reference (%) |
| --- | --- | --- | --- |
| 1 | RMRC 2 | 99 | 100 |
| 2 | RMRC 5 | 99 | 100 |
| 3 | RMRC 6 | 99 | 100 |
| 4 | RMRC 7 | 99 | 100 |
| 5 | RMRC 22 | 99 | 99 (M) |
| 6 | RMRC 23 | 99 | 100 |
| 7 | RMRC 24 | 99 | 100 |
| 8 | RMRC 27 | 99 | 100 |
| 9 | RMRC 28 | 99 | 100 |
| 10 | RMRC 30 | 99 | 100 |
| 11 | RMRC 38 | 99 | 100 |
| 12 | RMRC 46 | 99 | 100 |
| 13 | RMRC 90 | 99 | 100 |
| 14 | RMRC 103 | 99 | 100 |
| 15 | RMRC 104 | 100 | 100 |
| 16 | RMRC 106 | 99 | 100 |
| 17 | RMRC 108 | 99 | 100 |
| 18 | RMRC 110 | 99 | 100 |
| 19 | RMRC 112 | 99 | 100 |
| 20 | RMRC 154 | 99 | 100 |
| 21 | RMRC 155 | 99 | 100 |
| 22 | RMRC 156 | 99 | 100 |
| 23 | RMRC 157 | 100 | 100 |
| 24 | RMRC 158 | 100 | 100 |
| 25 | RMRC 159 | 99 | 100 |
| 26 | RMRC 160 | 99 | 100 |
| 27 | RMRC 162 | 99 | 100 |
| 28 | RMRC 163 | 99 | 100 |
| 29 | RMRC 164 | 99 | 100 |
| 30 | RMRC 165 | 99 | 100 |
| 31 | RMRC 166 | 99 | 100 |
| 32 | RMRC 167 | 99 | 100 |
| 33 | RMRC 168 | 99 | 100 |
| 34 | RMRC 170 | 99 | 100 |
| 35 | RMRC 171 | 99 | 100 |

### Sequence identification, alignment and phylogenetic analysis of RBD

The alignment of the spike sequences of RMRC isolates with the reference strain RBD, revealed the RBD region of Indian SARS-CoV-2 isolates sitting at 996-1578 bp region of the spike gene consisting of total 582 bases. The protein sequence of the RBD spans from 333 (T, Threonine) - 526 (G, Glycine) amino acids of the spike protein^21^. Interestingly RBD sequence alignment results showed that, RMRC 22 has 99 % sequence identity with the reference RBD and harbours a mutation at 1204 (Nucleotide: Ref A, RMRC 22 C; Protein: Ref I, RMRC 22 L) (Fig.1A). Except for RMRC 22, other three shared 100% identity with the reference RBD (Uniprot ID: PODTC2, 333-526 amino acids of the spike protein). As only RMRC 22 had a mutation, it was named as mutant isolate, whereas all other isolates were considered as wild type.

**Figure 1:**
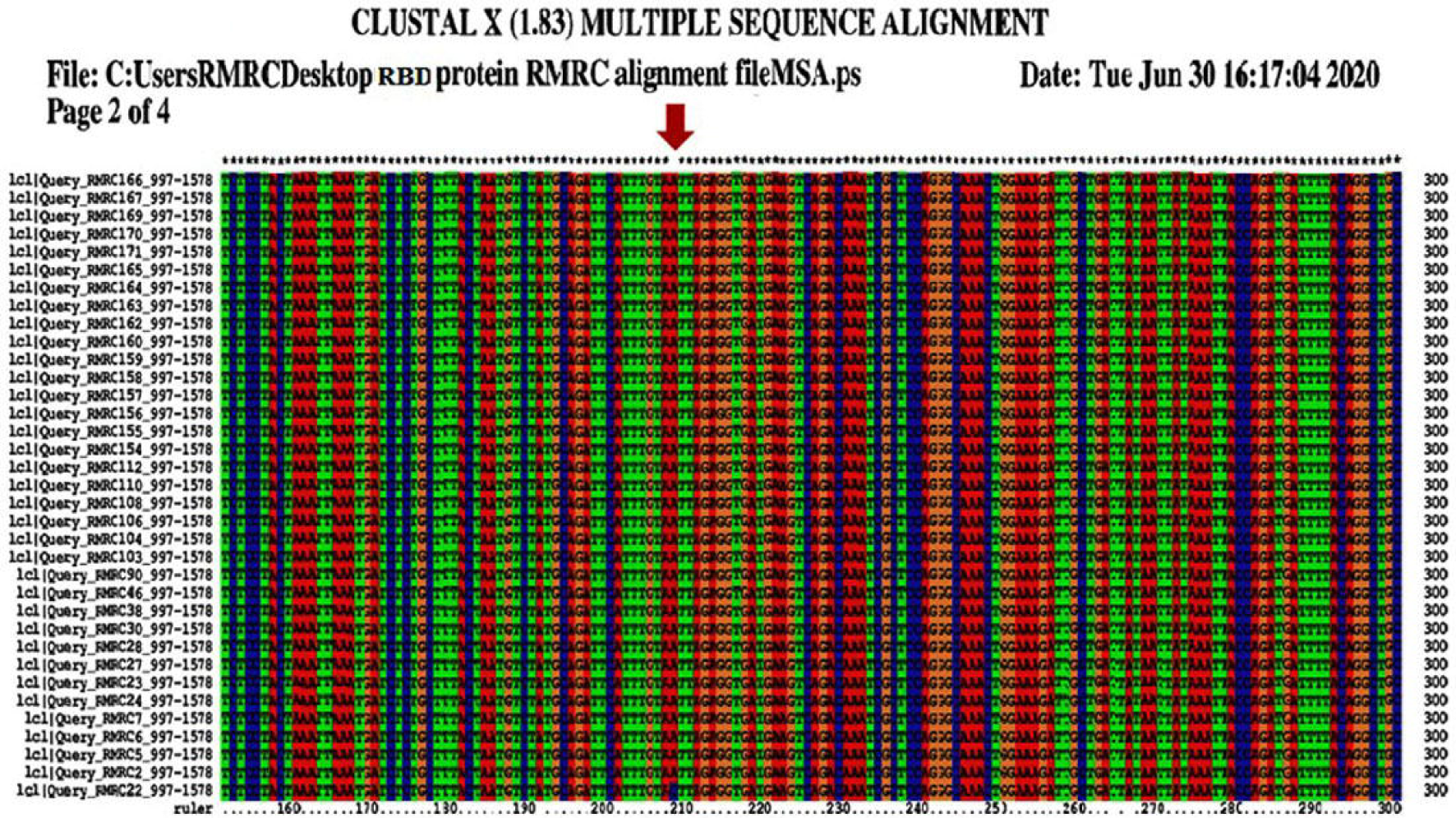
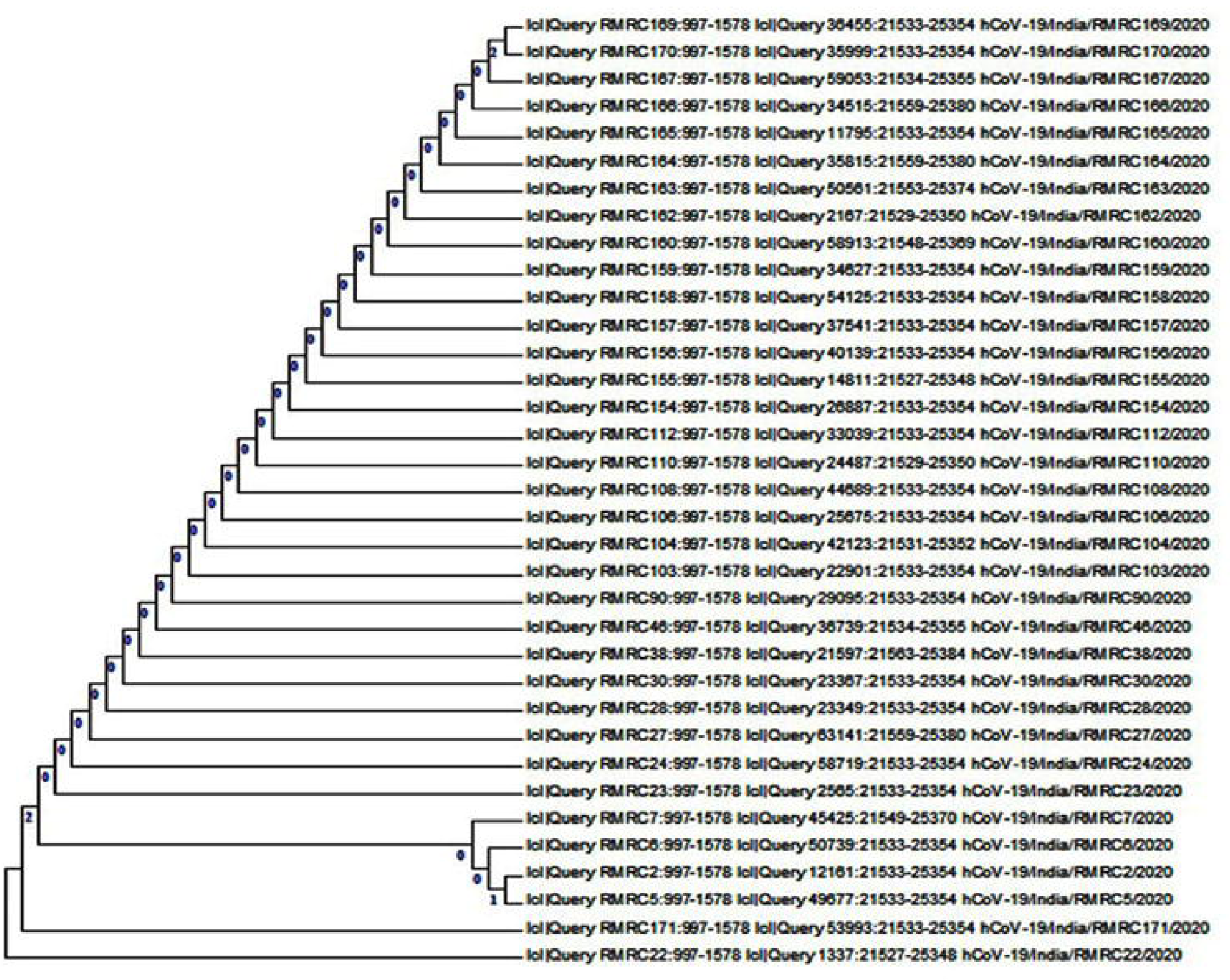
(A) Multiple sequence alignment and (B) phylogenetic analysis of RBD sequences of RMRC isolates.

The phylogenetic analysis was carried out using RBD sequences of all 35 RMRC isolates which showed that they formed four different clusters. RMRC 22 having a mutation belonged to first cluster, RMRC 171 RBD formed second cluster, four RBDs (RMRC 2, 5, 6 and 7) grouped in the third cluster and all other RBDs were in cluster four, which describes their phylogenetic distribution (Fig.1B).

### Protein structure analysis and interaction of RBD with ACE2 receptor

The sequence alignment analysis showed that ACE2 of pangolin, hamster, Chinese and Indian bat shared 85 %, 84 %, 80 % and 78 % identity with human ACE2 respectively (Suppl. Fig.1). The quality of modelled structures of ACE2 was validated using several computational methods (pangolin Fig.2A; hamster Fig.3A; Chinese bat Fig.4A; Indian bat Fig. 5A; human Fig.6A). In Pangolin, out of 805 amino acid residues, the Ramachandran analysis illustrated 637 (88.0%) residues in most favoured regions, 66 (9.1%) in additional allowed regions, 14 (1.9%) in generously allowed regions and 7 (1.0%) in disallowed regions (Fig. 2B). In Hamster, out of 785 amino acid residues, the Ramachandran analysis illustrated 613 (87.2%) residues in most favoured regions, 70 (10.0%) in additional allowed regions, 12 (1.7%) generously allowed regions and 8 (1.1%) in disallowed regions (Fig.3B). In Chinese bat, out of 805 amino acid residues, the Ramachandran analysis illustrated 634 (88.8%) residues in most favoured regions, 66 (9.2%) in additional allowed regions, 12 (1.7%) generously allowed regions and 2 (0.3%) in disallowed regions (Fig.4B). In Indian bat, out of 807 amino acid residues, the Ramachandran analysis illustrated 630 (86.8%88.8%) residues in most favoured regions, 73 (10.1%) in additional allowed regions, 15 (2.1%) generously allowed regions and 8 (1.1%) in disallowed regions (Fig.5B). Using the Qualitative Model Energy Analysis (QMEAN) server, the model quality was determined. The overall quality of model was good as indicated by its QMEAN Z-score and QMEAN4 global score. Low quality models are expected to have a negative QMEAN Z-score. The QMEAN4 ranges from 0 to 1 and a higher value indicates good qualitymodel^22^. Additionally, the overall quality of the model was evaluated using Protein Structure Analysis (ProSA) tool which provides a quality score (Z-score) as compared to all known protein structure from x-ray crystallography as well structural NMR. The obtained Z-score value was −6.33 (Pangolin), −4.76 (Hamster), −6.24 (Chinese bat) and −6.73 (Indian bat); which indicates the high quality of the models compared to known protein structures (Fig. 2B-5B).

**Figure 2:**
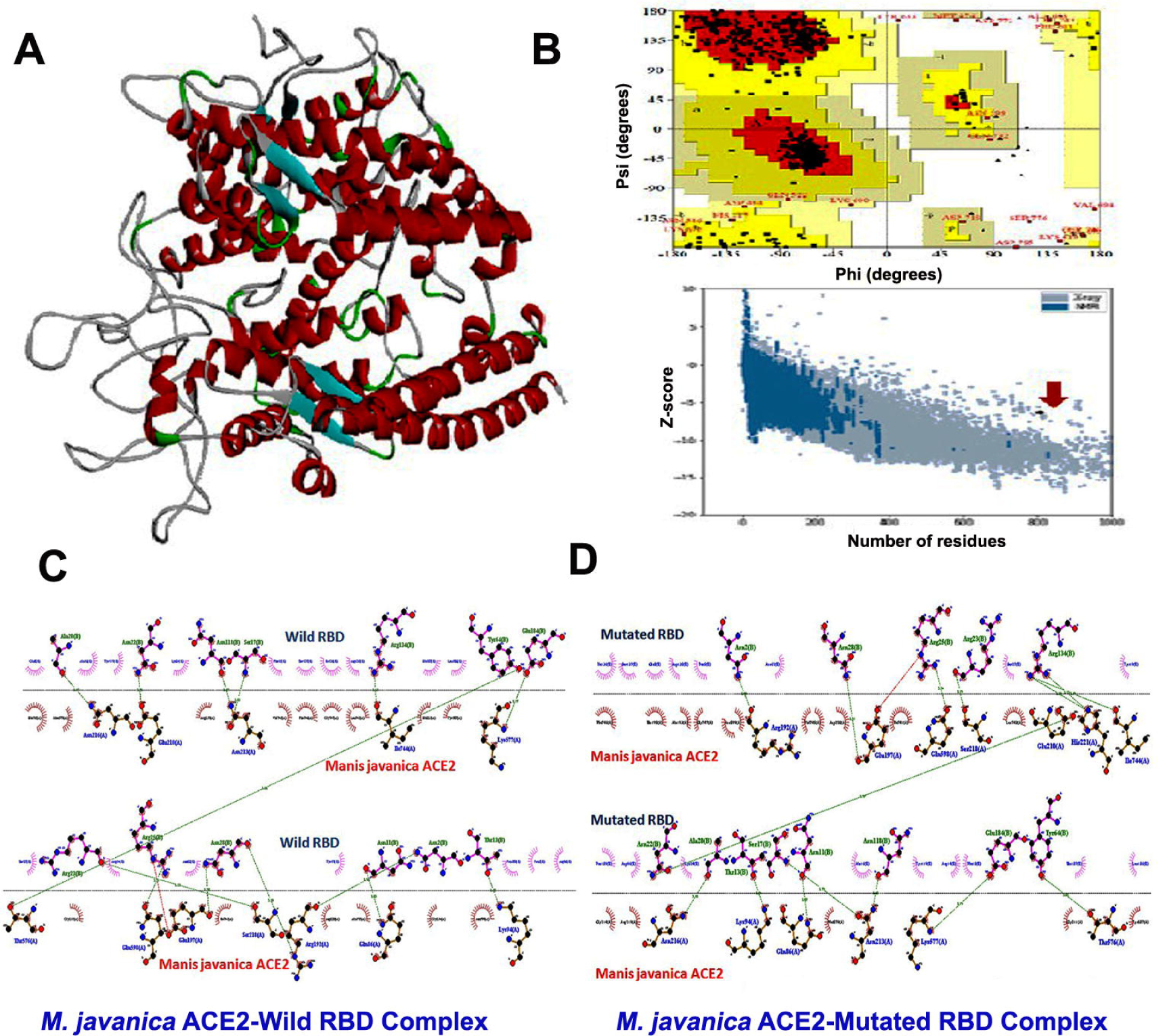
3D structure analysis and interaction of ACE2 receptor of *Manis javanica* with wild and mutated RBD. (A) Predicted 3D structure of ACE2 receptor; (B) Ramachandran plot and Z-score; (C) ACE2 interaction with wild type RBD and (D) mutated RBD.

**Figure 3:**
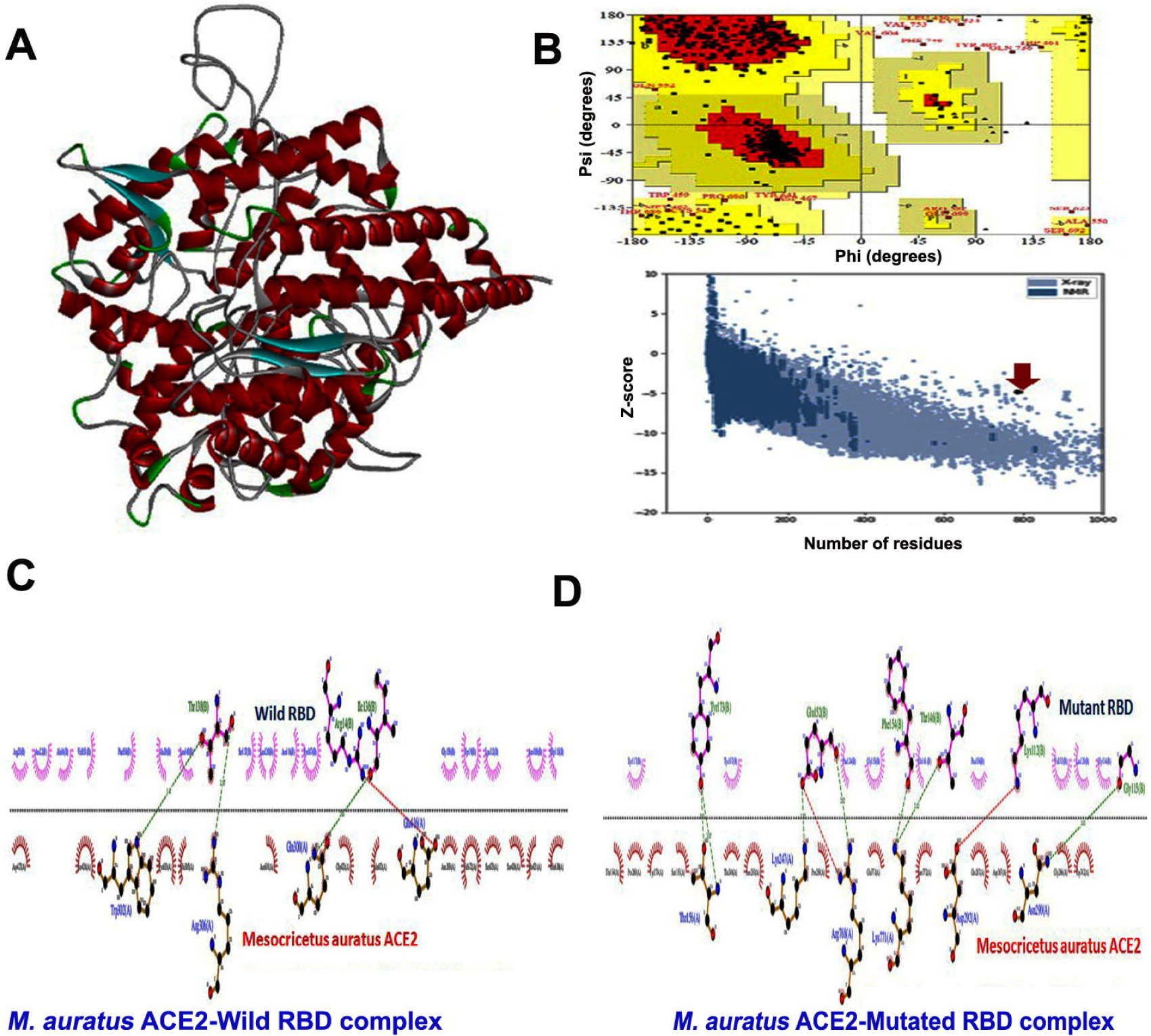
3D structure analysis and interaction of ACE2 receptor of *Mesocricetus auratus* with wild and mutated RBD. (A) Predicted 3D structure of ACE2 receptor; (B) Ramachandran plot and Z-score; (C) ACE2 interaction with wild type RBD and (D) mutated RBD.

**Figure 4:**
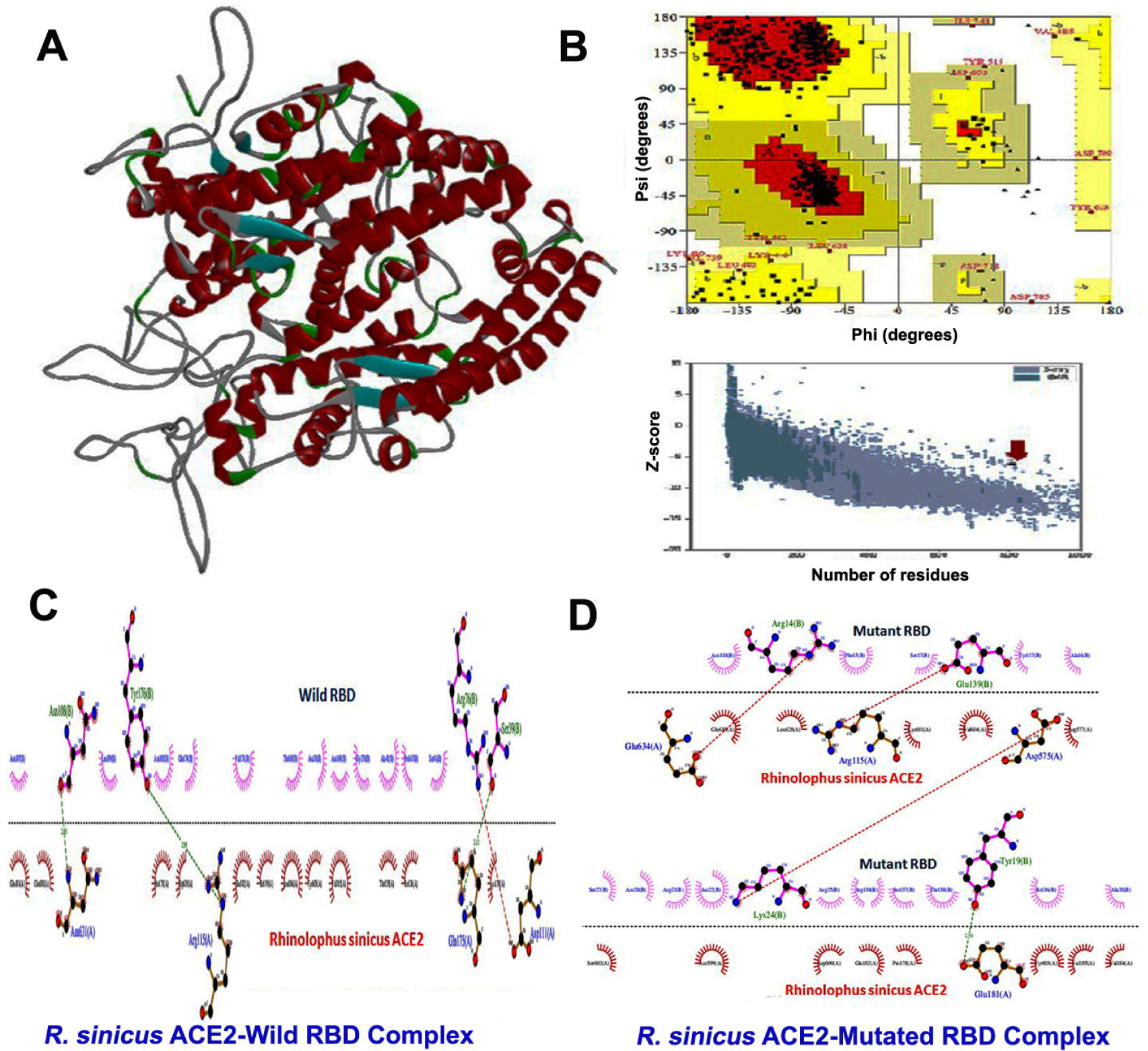
3D structure analysis and interaction of ACE2 receptor of *Rhinolophus sinicus* with wild and mutated RBD. (A) Predicted 3D structure of ACE2 receptor; (B) Ramachandran plot and Z-score; (C) ACE2 interaction with wild type RBD and (D) mutated RBD.

**Figure 5:**
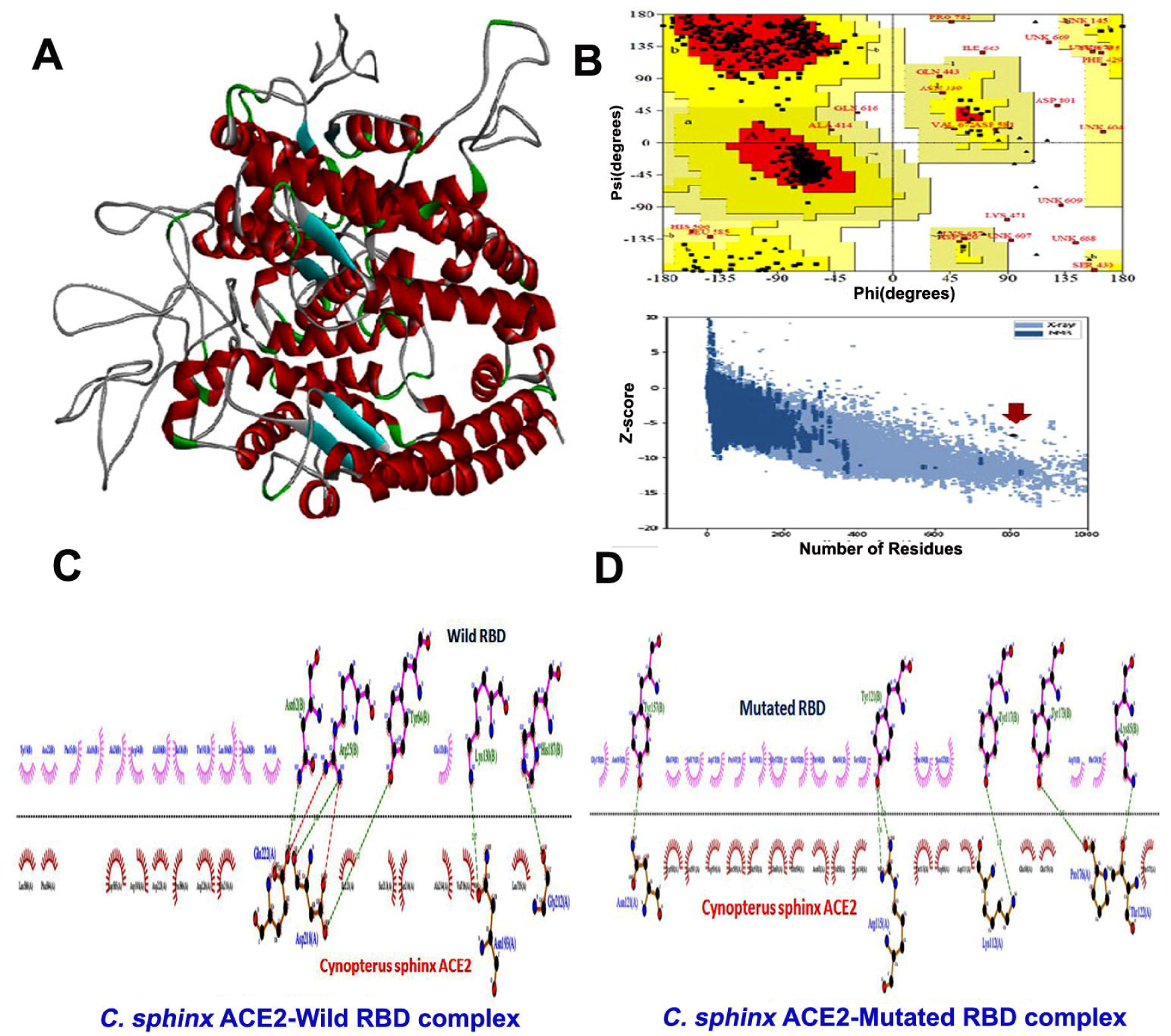
3D structure analysis and interaction of ACE2 receptor of *Cynopterus sphinx* with wild and mutated RBD. (A) Predicted 3D structure of ACE2 receptor; (B) Ramachandran plot and Z-score; (C) ACE2 interaction with wild type RBD and (D) mutated RBD.

**Figure 6:**
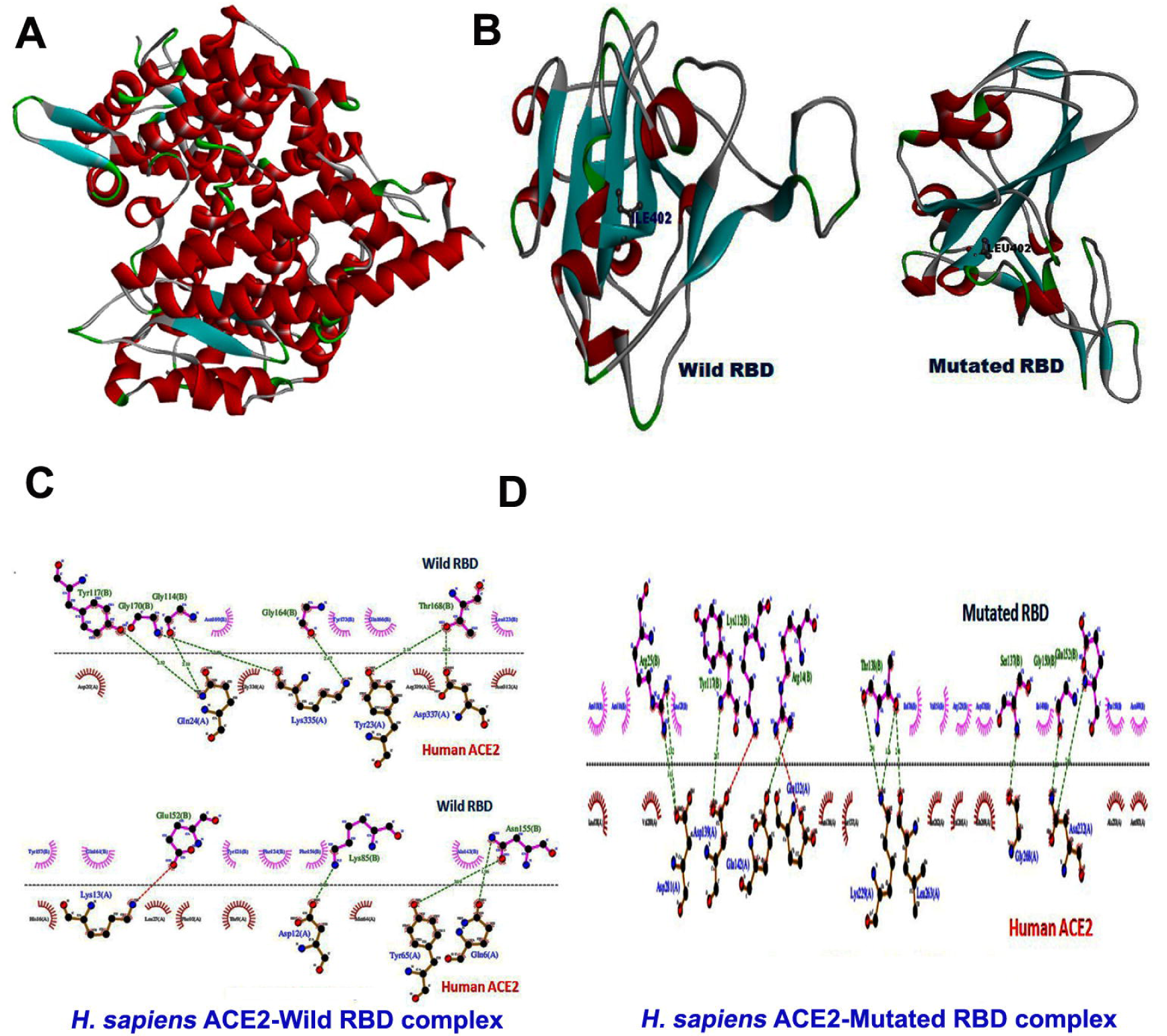
3D structure and interaction of ACE2 receptor of *Homo Sapiens* with wild and mutated RBD. (A) 3D structure of ACE2 receptor; (B) predicted structure of wild and mutated RBD of RMRC SARS-CoV-2 isolates; (C) *H. sapiens* ACE2 interaction with wild type RBD and (D) mutated RBD.

As the 3D structure of human RBD was already available in the database (PDB ID-6M0J), we inserted the observed mutation of RMRC 22 isolate (Protein: Ref I, RMRC 22 L) and predicted the structure of mutated RBD (Fig. 6B). As the specific binding of viral RBD with host ACE2 receptor determines the establishment of infection, we analysed the interaction of Indian SARS-CoV-2 RBDs (both mutant and wild type) with the ACE2 receptor of human as well as other species which are reported to be natural reservoirs for this virus. The interaction analysis showed that, the mutated RBD of RMRC 22 isolate has stronger interaction with human ACE2 (Fig. 6D) as compared to the wild type RBD (Fig. 6C). The interaction between mutated RBD-human ACE2 has a binding energy of **-**65.95 Kcal/Mol with six hydrogen bonds whereas wild type RBD-human ACE2 interaction has -63.09 Kcal/Mol binding energy with four hydrogen bonds. For ACE2 receptor of all other species (Pangolin: Fig. 2C & 2D; hamster: Fig.3C & 3D; Chinese bat: Fig.4C & 4D; Indian bat: Fig.5C & 5D), wild type RBD seems to have stronger binding affinity with no difference in number of hydrogen bonds (except hamster, wild type RBD-ACE2:2 and mutant RBD-ACE2: 3 hydrogen bonds). The details of the interaction analysis with H-bond forming residues and average distance of H-bonds has been described in Table-2. However, if we compare the interaction of mutated and wild type RBD with ACE2 receptor of all species, the interaction between mutated RBD and human ACE2 is the strongest one with highest binding energy and highest number of hydrogen bonds.

**Table 2:** Protein–protein interaction analysis of ACE2 and RBD (wild and Mutated) obtained by HawkDock web server.

| Sl. No . | Receptor PDB | Ligand PDB | Binding Energy(Kcal/Mol) | No. Of H -Bonds | H-Bond Forming Residues | Average Distance Of H-Bonds (Å) |
| --- | --- | --- | --- | --- | --- | --- |
| 1. | Human ACE2 | Wild RDB | -63.09 | 4 | GLN24,TYR65,LYS335,GLU17,LYS335 | 2.833 |
| 2. | Human ACE2 | Mutated RDB | -65.95 | 6 | LYS229,ASN232,ASP139,GLY268,LEU263, ASN232 | 2.853 |
| 3. | Mesocricetus auratus (Hamster) | Wild RDB | -56.75 | 2 | ARG306, ILE421 | 2.709 |
| 4. | Mesocricetus auratus (Hamster) | Mutated RDB | -50.4 | 3 | THR156:N , THR156, ASN290 | 3.141 |
| 5. | Rhinolophus sinicus (Chinese Bat) | Wild RDB | -51.81 | 4 | ASN631,GLU181,ASN631, GLU182 | 2.797 |
| 6. | Rhinolophus sinicus (Chinese Bat) | Mutated RDB | -27.09 | 4 | ARG577,ARG577,GLU181, ASN599 | 3.036 |
| 7. | Cynopterus sphinx (Indian Bat) | Wild RDB | -37.53 | 4 | ASP218,GLU222, GLU222, ASN193 | 2.667 |
| 8. | Cynopterus sphinx (Indian Bat) | Mutated RDB | -23.34 | 4 | LYS112, ASN121,THR122,PRO176 | 3.024 |
| **9.** | **Manis**<br>**javanica**<br>**(Pangolin)** | **Wild RDB** | **-70.92** | **13** | **A:GLN86,LYS94,ASN213,ASN216,SER218**<br>**,THR576,ARG192,ASN213,GLY214,GLU2**<br>**10,GLN598,GLU197,ILE744** | **2.807** |
| **10**<br>**.** | **Manis**<br>**javanica**<br>**(Pangolin)** | **Mutated**<br>**RDB** | **-72.74** | **11** | **GLN86,LYS94, ASN213, ASN216,THR576,**<br>**ARG192, GLY214, GLU210,GLN598,**<br>**GLU197,ILE744** | **2.860** |

## 4. Discussion

The complex trajectory of the recent COVID-19 pandemic in India poses greater risk towards control and containment of the infection. It is high time to understand its mobility pattern in the country and the viral genetic properties favouring its virulence. Though during early phase of pandemic, Odisha, had comparatively very few positive cases as well as small number of deaths, gradually, the virus has become rapidly infectious making the clinical scenario worsen. Many people from Odisha were working outside the state and during this pandemic, they returned to their home state due to many reasons. The samples included in our study were mainly collected from suspected cases who had a travel history from different states with one foreign travel record.

As spike protein of SARS-CoV-2 mediates viral entry in to host and houses RBD, which binds to ACE2 receptor of host cell, understanding the spike-RBD distribution in the genome of Indian isolates is crucial for therapeutic design. SARS-CoV-2 spike protein is reported to have a stronger binding affinity for ACE2 than SARS-CoV and higher affinity means low number of virus is required to infect the cell, which may explain the high transmission of SARS-CoV-2^23^. In the current study, spike region of 32 isolates showed altered nucleotide bases at multiple positions as compared to the Wuhan reference strain suggesting mutations in these Indian isolates during the spread. For SARS-CoV-2, specific RBD-ACE2 binding ensures infection as well as serves as a potential target for developing treatment strategies for this infection^23^. According to Premkumar et al. 2020, the RBD of SARS-CoV-2 is an immunodominant and a potential target of antibodies in COVID-19 patients^24^. The mutation found in RMRC 22 isolate might play a role in altering the antigenicity or binding affinity of the respective RBD. Due to the rapid spreading and evolution, SARS-CoV-2 RBD is known to acquire several mutations leading to increased binding affinity to human ACE2 receptor^25^. In France, multiple mutations were identified in RBD of SARS-CoV-2 contributing to higher receptor binding capacity, which might be responsible for increased virus spread and infectivity^26^. On the other hand, a mutation in S protein has been found to be associated with decrease in receptor binding affinity^27,28^. In the current study, mutation in the RBD region of Indian isolates did not seem to affect its interaction with ACE2 receptor of other species prominently except that of human. Surprisingly, the patient, from whom, the RMRC 22 isolate was obtained, had a travel history of returning from Nizamuddin (cluster detected from New Delhi, India during April, 2020) recently before he tested positive for SARS-CoV-2. However, no mutation was observed in the isolates obtained from his other family members (his father and two brothers) who were also COVID-19 positive. It appears that, emergence and role of a mutation in any region of SARS-CoV-2 genome depends upon multiple factors including the geographical distribution, rate of spreading, alteration in the virulence of the virus and immune response of the host. The interaction analysis of mutated and wild type RBDs with ACE2 receptor indicated that bats and pangolins could be suitable natural reservoirs for Indian isolates of this virus and this finding falls parallel with other earlier reports^29,30^. As hamster has been reported as a suitable animal model to study SARS-CoV-2 pathogenesis, the interaction of RBD of the Indian isolates included in the current study makes the earlier report more relevant^10^. Though the susceptibility of infection and death rate could be affected by several factors, mutation in the virus genome and its ability to adapt to new environment could be crucial.

Being an important determinant in SARS-CoV-2 infection, RBD-ACE2 interaction has already become the potential target for developing treatment therapy against this deadly pathogen. The current study provides important information regarding the structural basis of spike and RBD regions of few Indian SARS-CoV-2 isolates which gives an idea about their evolution and spreading. The mutation observed in the RBD region of one of the isolates sheds light on drug targeted therapy for different strains of virus. Further studies with larger number of isolates of a more wide origin would be helpful to understand this mutation pattern in RBD of Indian isolates.

## Supporting information

Supplementary figure: Sequence alignment of ACE2 receptors of Manis javanica, Mesocricetus auratus, Rhinolophus sinicus, Cynopterus sphinx and Homo Sa

## Acknowledgment

We acknowledge the contribution of all the VRDL staff (Laboratory technicians, Multi tasking workers) whose relentless effort in COVID testing has given the strength to further plan up new experiments and studies. We also acknowledge the support of Dr. L.M. Ho for his contribution. We acknowledge Dr. Ira Praharaj and Dr. Sidharth Giri for their expert advice and scientific inputs.

**Supplementary figure:** Sequence alignment of ACE2 receptors of *Manis javanica, Mesocricetus auratus, Rhinolophus sinicus, Cynopterus sphinx and Homo Sapiens.*

