## Supplementary figures and images for "Sequence analysis of Indian SARS-CoV-2 isolates shows a stronger interaction of mutated receptor binding domain with ACE2 receptor"

### Supplementary figure: Sequence alignment of ACE2 receptors of Manis javanica, Mesocricetus auratus, Rhinolophus sinicus, Cynopterus sphinx and Homo Sa

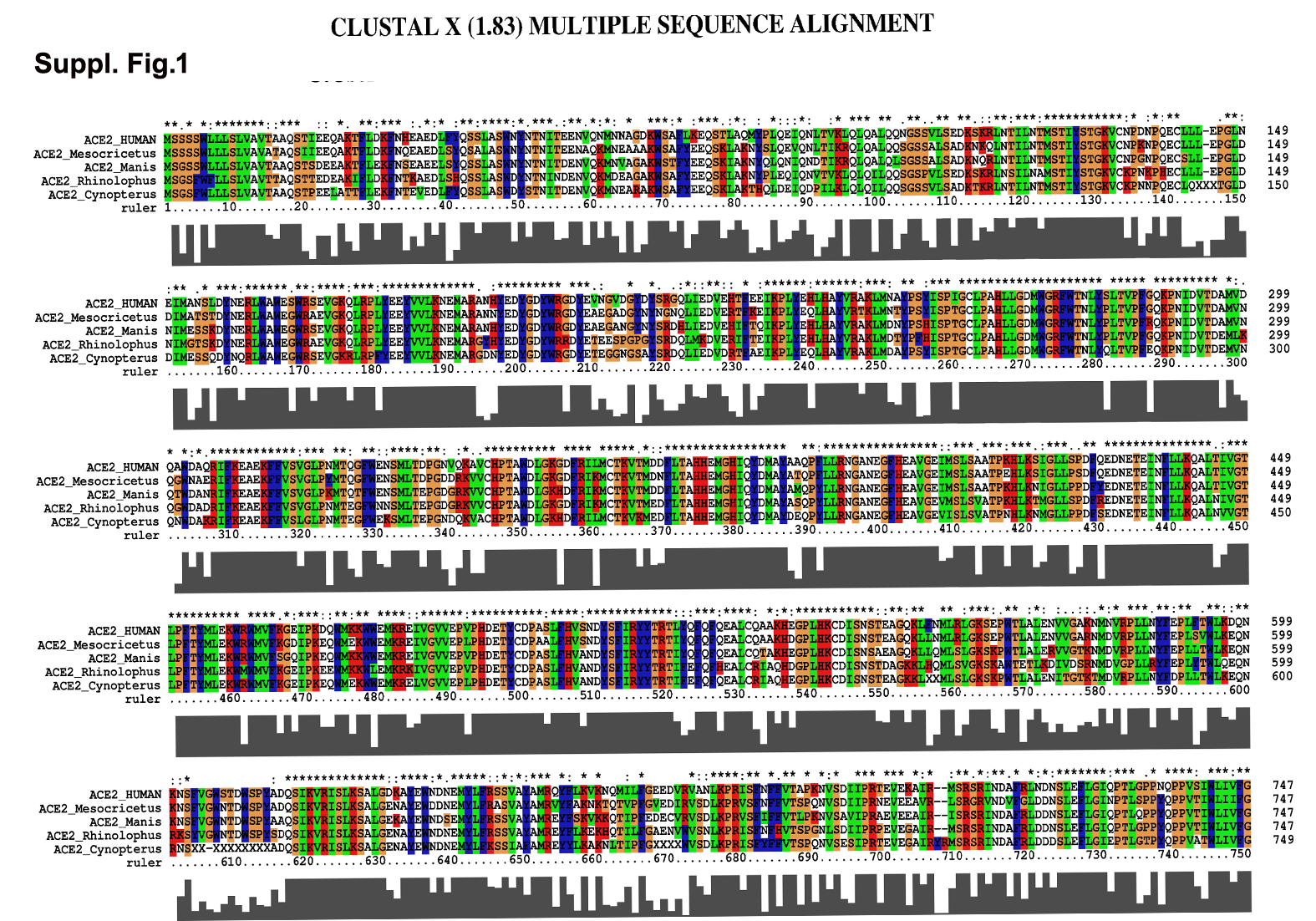
